# A *De Novo* regulation design shows an effectiveness in altering plant secondary metabolism

**DOI:** 10.1101/2020.12.20.423678

**Authors:** Mingzhuo Li, Xianzhi He, Christophe La Hovary, Yue Zhu, Yilun Dong, Shibiao Liu, Hucheng Xing, Yajun Liu, Yucheng Jie, Dongming Ma, Seyit Yuzuak, De-Yu Xie

## Abstract

**Introduction:** Transcription factors (TFs) and cis-regulatory elements (CREs) control gene transcripts involved in various biological processes. We hypothesize that TFs and CREs can be effective molecular tools for *De Novo* regulation designs to engineer plants.

**Objectives:** We selected two Arabidopsis TF types and two tobacco CRE types to design a *De Novo* regulation and evaluated its effectiveness in plant engineering.

**Methods:** G-box and MYB recognition elements (MREs) were identified in four *Nicotiana tabacum JAZs* (*NtJAZs*) promoters. MRE-like and G-box like elements were identified in one nicotine pathway gene promoter. TF screening led to select Arabidopsis Production of Anthocyanin Pigment 1 (PAP1/MYB) and Transparent Testa 8 (TT8/bHLH). Two *NtJAZ* and two nicotine pathway gene promoters were cloned from commercial Narrow Leaf Madole (NL) and KY171 (KY) tobacco cultivars. Electrophoretic mobility shift assay (EMSA), cross-linked chromatin immunoprecipitation (ChIP), and dual luciferase assays were performed to test the promoter binding and activation by PAP1 (P), TT8 (T), PAP1/TT8 together, and the PAP1/TT8/Transparent Testa Glabra 1 (TTG1) complex. A DNA cassette was designed and then synthesized for stacking and expressing PAP1 and TT8 together. Three years of field trials were performed by following industrial and GMO protocols. Gene expression and metabolic profiling were completed to characterize plant secondary metabolism.

**Results:** PAP1, TT8, PAP1/TT8, and the PAP1/TT8/TTG1 complex bound to and activated *NtJAZ* promoters but did not bind to nicotine pathway gene promoters. The engineered red P+T plants significantly upregulated four *NtJAZs* but downregulated the tobacco alkaloid biosynthesis. Field trials showed significant reduction of five tobacco alkaloids and four carcinogenic tobacco specific nitrosamines in most or all cured leaves of engineered P+T and PAP1 genotypes.

**Conclusion:** G-boxes, MREs, and two TF types are appropriate molecular tools for a *De Novo* regulation design to create a novel distant-pathway cross regulation for altering plant secondary metabolism.

## Introduction

Plant MYB and bHLH proteins are two large TF families with diverse regulations involved in various biological processes [1–4]. To date, TFs that regulate the anthocyanin biosynthesis have gained a comprehensive characterization in both model and non-model plants. MYB and bHLH TFs together with a WD40 protein have been demonstrated to form MBW complexes to regulate the biosynthesis of anthocyanins [5–7]. In particular, the MBW complex formed by Arabidopsis Production of Anthocyanin Pigment 1 (PAP1), Transparent Testa 8 (TT8), and Transparent Testa Glabra 1 (TTG1) has been characterized to be a main regulatory complex. *PAP1*, *TT8*, and *TTG1* encode a R2R3-MYB TF (namely MYB75) [8], bHLH TF (namely bHLH42) [9], and a WD40 protein [10], respectively. Moreover, the functions of these three genes in Arabidopsis and their homologs in other plants have been intensively characterized in genetics and biochemistry [11]. PAP1 and TT8 bind to MYB recognizing elements (MREs) and G-box elements in promoters of anthocyanin pathway genes (such as *DFR*, *ANS* and *3-GT*, Fig. 1 A), respectively [12–14]. The current dogma is that on the one hand, PAP1, TT8, and TTG1 form a master regulatory MYB-bHLH-WD40 (MBW) complex to only activate the anthocyanin biosynthesis in plants (Fig. 1 A) [6, 15–17]; on the other hand, Arabidopsis JAZ repressors can interact with PAP1, TT8, and their MBW complex to negatively regulate the anthocyanin biosynthesis in the absence of jasmonate (JA) [18, 19].

**Figure 1.**
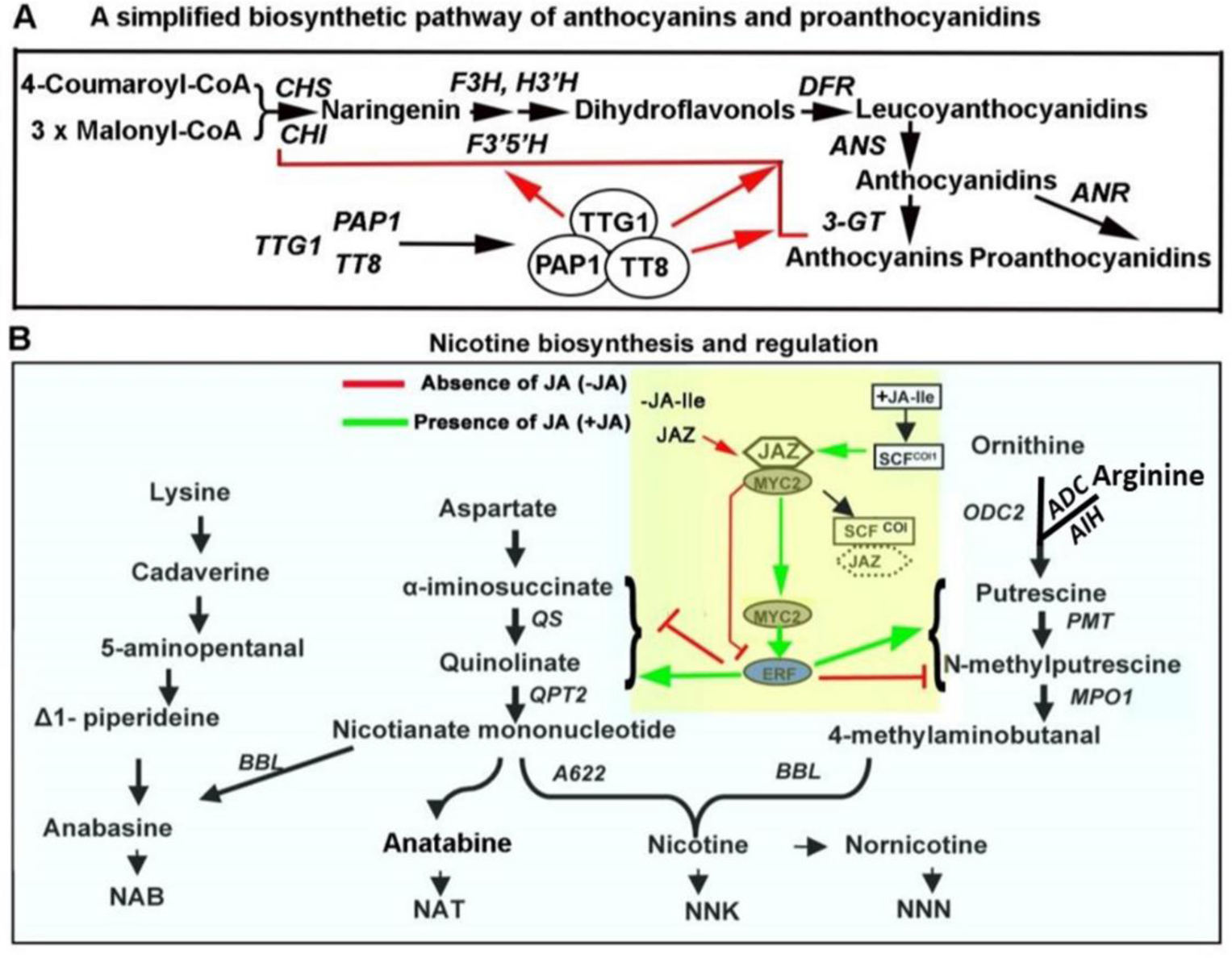
Biosynthetic pathways of nicotine, other tobacco alkaloids, and anthocyanin and their regulations. **A**, the biosynthetic pathway of Arabidopsis anthocyanin from the step catalyzed by CHS. *PAP1*, *TT8*, and *TTG1*encode MYB75, bHLH, and WD40 to form a MYB-bHLH-WD40 complex that activates late pathway genes. Gene abbreviations are *CHS* and *CHI*: chalcone synthase and isomerase, *F3H*: flavonone-3-hydroxylase, *F3’H* and *F3’5’H*: flavonoid-3’-hydroxylase and 3’5’hydroxylase, *DFR*: dihydroflavonol reductase, *ANS*: anthocyanidin synthase, *3- GT*: glycosyltransferase. *ANR* encodes anthocyanidin reductase, a key enzyme toward proanthocyanidin biosynthesis. **B,** the biosynthetic pathway of nicotine, nornicotine, and tobacco specific nitrosamines (TSNAs) and the regulation mechanism. MYC2 and EAR are two positive transcription factors that activate the expression of most pathway genes. In the absence of isoleucine-jasmonate (Ile-JA), NtJAZ1 and NtJAZ3 repressors bind to MYC2 to inhibit the activation of the pathway genes (red lines). In the presence of bioactive Ile-JA (green arrows), SCF^COI1^ perceives this plant hormone to form a new JA-SCF^COI1^ complex to bind to NtJAZs and lead them to proteasomal ubiquitination. The degradation of NtJAZs releases MYC2 to activate pathway gene expression toward the biosynthesis of nicotine. Gene abbreviations are *AO*: aspartate oxidase, *QS*: quinolinate synthase, *QPT2*: quinolinate phosphoribosyltransferase 2, *ADC:* arginine decarboxylase, *AIH*: agmatine iminohydrolase, *ODC*: ornithine decarboxylase, *MPT1*: N-methylputricine transferase 1, *MPO1*: methylputricine oxidase 1, *A622*: an isoflavone reductase like enzyme, *BBL*: berberine bridge enzyme- like protein, and *NND*: nicotine N-demethylase.

Moreover, *PAP1* has been used to create red crop varieties to engineer anthocyanins, such as tobacco [20, 21], tomato [22, 23], hop [24], and canola [25]. All these studies reported that PAP1 alone could activate the anthocyanin biosynthesis in different tested plants. However, whether PAP1, its partner TT8, and their homologs can regulate plant secondary metabolism remains open for studies.

Tobacco is an ideal model plant to study plant secondary metabolism, given that it produces thousands of metabolites in the alkaloid, phenolic, and terpenoid families [21, 26–28]. Nicotine is the main alkaloid of tobacco (*Nicotiana tabacum*) and harmfully addictive [29–36]. Other harmful tobacco alkaloids include nornicotine, anabasine, anatabine, and myosmine (Fig. 1 B). Tobacco specific nitrosamines (TSNAs) are carcinogenic compounds in tobacco smoke. Five common TSNAs are N-nitrosonornicotine (NNN), nicotine-derived nitrosamine ketone (NNK), N’-nitrosoanatabine (NAT), N-nitrosoanabasine (NAB), and 4-(methylnitrosamino)-1-(3- pyridyl)-1-butanol (NNAL) [37–39]. Given that NNN is a severe carcinogen, the Food and Drug Administration has proposed a guideline that the average level of NNN must be less than 1.0 ppm (dry weight) in all final smokeless tobacco products [40].

To reduce these harmful compounds, numerous achievements have been accomplished in elucidating the biosynthesis of nicotine and other tobacco alkaloids [41–47]. Nicotine is biosynthesized from two distinct pathways (Fig. 1 B), the steps of which are catalyzed by enzymes encoded by 10 genes *AO*, *QS*, *QPT2*, *ODC*, *ADC*, AIH, *PMT*, *MPO*, *A622* and *BBL*, (Fig. 1 B) [42, 48, 49]. In addition, *CYP82E4* and its homologs have been characterized to involve the demethylation of nicotine to form nornicotine [50–52]. The mechanisms of the anatabine and anabasine formation are unknown [42]. One hypothesis is that these two alkaloids result from the BBL catalysis of nicotinic acid. The other is that anabasine is derived from lysine, a different pathway (Fig. 1 B) [42]. Four main TSNAs, NNK, NNN, NAT, and NAB (Fig. 1 B), result from the nitrosation of nicotine, nornicotine, anatabine, and anabasine during the leaf curing process, respectively [42]. Meanwhile, the regulation of the nicotine pathway has gained an intensive elucidation. Myelocytomatosis oncogene (MYC2) and APETALA 2/ethylene response factor (AP2/ERF) of tobacco are two groups of transcription factors (TFs) that directly bind to promoters of nicotine pathway genes [42, 48, 53–55]. Moreover, jasmonate (JA), a plant hormone, essentially controls the nicotine biosynthesis (Fig. 1B) [56–59]. JA regulates the nicotine biosynthesis via a co-receptor complex consisting of coronatine insensitive 1 (COI1), a Skp/Cullin/F-box (SCF^COI1^) complex, and JA ZIM-domain (JAZ) repressor [53–55, 60]. The tobacco genome has been reported to have 15 NtJAZs, in which NtJAZ1, NtJAZ3, NtJAZ7, and NtJAZ10 are associated with the regulation of the nicotine biosynthesis [54, 60, 61]. When tobacco roots lack isoleucine-JA (JA-Ile), NtJAZ1 and NtJAZ3 bind to NtMYC2 and then turn off the nicotine biosynthesis; when JA-Ile exists, it induces the interaction of SCF^COI1^ and JAZ to form a complex, which leads to the ubiquitination of JAZs to release NtMYC2. The freed NtMYC2 then activates the biosynthetic pathway of nicotine (Fig. 1 B) [54, 55, 60]. Compared with the appropriate characterization of NtJAZs’ functions, how their transcription in tobacco is regulated remains open for investigations. To date, based on those pathway and regulatory genes, on the one hand, antisense and RNAi have been reported to down-regulate gene expression (such as *BBL*, *ODC*, *QPT*, and nicotine demethylase gene) to reduce nicotine, nornicotine, or TSNAs in leaves [49, 55, 62, 63]. On the other hand, these multiple successes revealed a challenge that the reduction of nicotine was traded with the increase of anatabine and other alkaloids [64, 65]. These successful engineering data indicate that continuous efforts are necessary to diminish TSNAs, nicotine, and other alkaloids simultaneously.

Since 2000 when more than 1500 TFs were annotated in the genome of *Arabidopsis thaliana* in [66], 320,370 TFs had been identified from 156 sequenced plant genomes and sequences of nine additional plants by 2017 [67]. Likewise, 21,997,501 TF bind sites (motifs or regulatory elements) (TFBSs) were determined in 63 plant genomes and nearly 2,493,577 TFBSs (cis-regulatory elements) were localized in the promoters [68]. Given that the interaction of TFs and TFBSs regulates plant development, growth, responses to stresses, and metabolisms, we hypothesize that the large numbers of cis-regulatory elements and TFs are useful molecular tools for *de novo* designs to create novel regulations for plant engineering. For proof of concept, herein, we selected tobacco crop to test our hypothesis, given that as described above, the success might enhance diminishing harmful nicotine, other tobacco alkaloids (OTAs), and TSNAs for human health. As introduced, given that *NtJAZ1* and *NtJAZ3* encode two main repressors of the nicotine biosynthesis, we hypothesized that their cis- regulatory elements were appropriate molecular tools to screen other plant TFs for *de novo* regulation designs and positive designs could be tested in the downregulation of nicotine biosynthesis in tobacco plants. Then, we analyzed cis-regulatory elements of all *NtJAZ* promoters and cloned MRE(s) and/or G-box(es) from four *NtJAZ* genes. In addition, we analyzed and identified MRE-like and G-box like elements of one nicotine pathway gene. Meanwhile, we analyzed Arabidopsis TFs that have a regulatory function to bind to these types of elements. As a result, PAP1 and TT8, two master regulators of Arabidopsis anthocyanin biosynthesis, were identified for this design. Mechanistic, transgenic, transcriptional, and metabolic quantification, and field farming practices showed that this design was successful to create a distant pathway-cross regulation (DPCR) crossing from the anthocyanin to nicotine pathway. The constructed DPCR showed an effectiveness in significantly reducing nicotine, four OTAs, and four TSNAs in tobacco plants, simultaneously. Moreover, the resulting data disclose new regulatory functions of both PAP1 and TT8 and new activators of four *NtJAZs*.

## Materials and methods

### Tobacco varieties

Six genotypic tobacco (*Nicotiana tabacum*) cultivars were used in this study. Two commercial dark tobacco varieties were Narrow Leaf Madole (NL) and KY171 for smokeless products (Andersen et al., 1990; Pearce et al., 2015). Two red transgenic genotypes were generated from NL and KY171 described below. Two others were *N. tabacum* Xanthi (an oriental tobacco variety) and a red PAP1 tobacco variety that is a novel homozygous red Xanthi variety (Fig. S22 B) created from the overexpression of the *Arabidopsis PAP1* (MYB75) (Xie et al., 2006). *Nicotiana benthamiana* was grown for dual-luciferase assay. Growth conditions in the greenhouse and phytotron are listed in supporting materials.

### Cloning of NtJAZ1, NTJAZ3, PMT2, and ODC2 promoters from Narrow Leaf Madole and KY171 cultivars

The promoter sequences of tobacco *NtJAZ1, NtJAZ3, NtJAZ7, NtJAZ10, PMT2*, and *ODC2* were identified from the genomic sequences of the NT90 cultivar curated at NCBI (https://www.ncbi.nlm.nih.gov/). The sequences identified were 1000 bp or 1500 bp. Based on these sequences, primers (Table S1) were designed for PCR to amplify *NtJAZ1, NtJAZ3, PMT2*, and *ODC2* promoters from the genomic DNA of wild type NL and KY171 tobacco plants, and then the amplified genomic DNA fragments were cloned to pEASY-T1 for sequencing. Briefly, the DNA fragments from PCR were purified using a gel-extraction Kit (Thermo Fisher, Waltham, MA) by following the manufacturer’s instruction and then ligated to the pEASY-T1 plasmid using a T4-ligase system (10 µl reaction system: 1 µl T4 buffer, 1 µl T4-ligase, 1.0 µl pEASY-T1 linear plasmid and 7.0 µl purified PCR product). The resulting recombinant plasmid was transformed into the competent *E. coli* Top10 cells, which were screened on agar-solidified LB medium supplemented with 50 mg/l ampicillin. Positive colonies were selected for sequencing at Eton Bio (Durham, NC, USA). All promoter sequences obtained were 1000 or 1500 bp (Figs. S1, S2, and S4). Analysis of promoter sequences and identification of regulatory elements such as AC- and AC-like elements (Figs. S1, S2, and S4) were completed with the PLACE (http://www.dna.affrc.go.jp/PLACE/) and PlantPan (http://plantpan.itps.ncku.edu.tw/) tools. The *NtJAZ1, NtJAZ3, PMT2*, and *ODC2* promoters were used for electrophoretic mobility shift assay, dual-luciferase, and ChIP assays described below.

**Figure 2.**
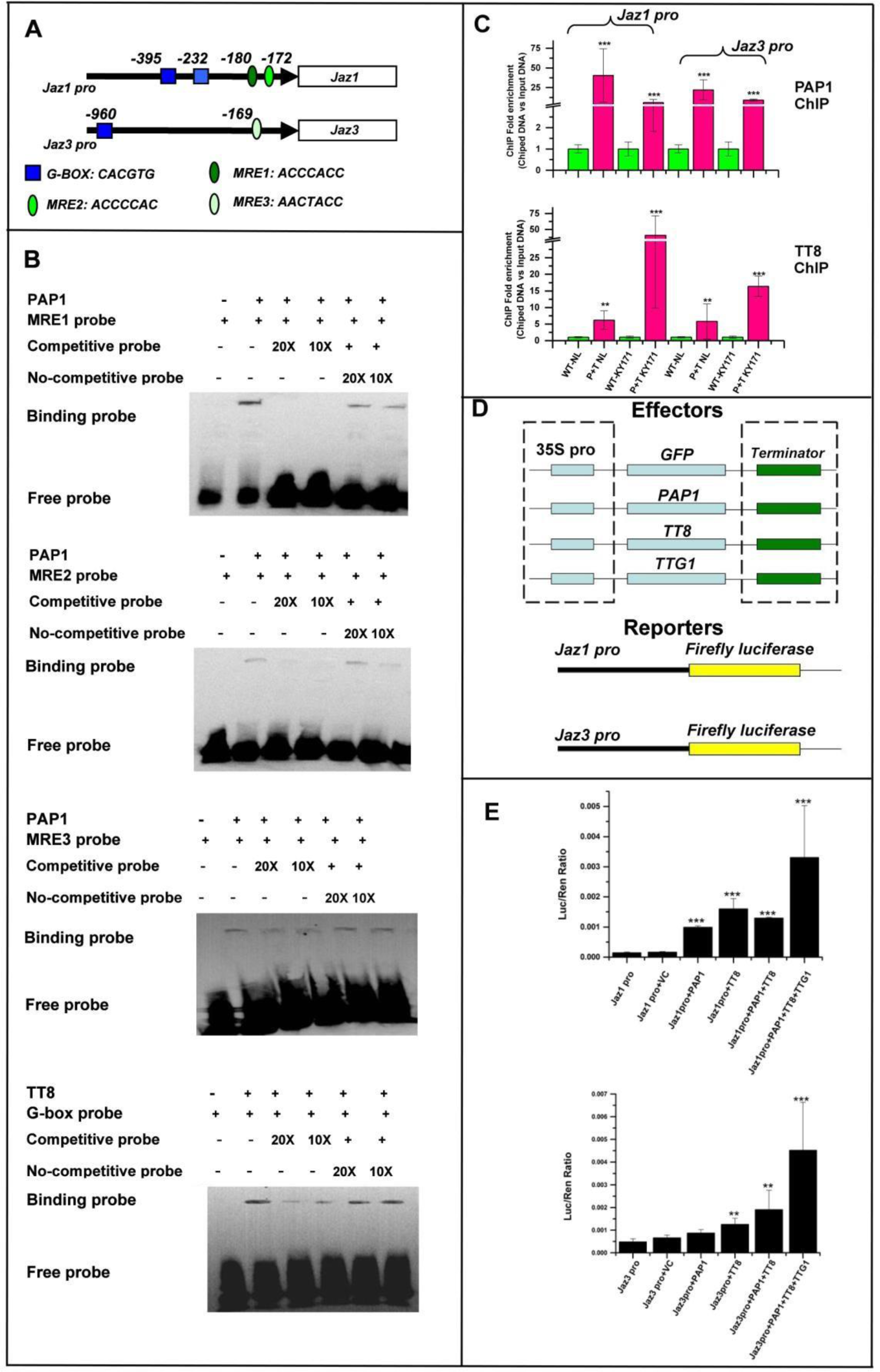
Binding and activation of *NtJAZ1* and *NtJAZ3* promoters by PAP1 and TT8 alone, PAP1 and TT8 together, and PAP1-TT8-WD40 complex. A, three MYB response element (MRE) types and G-box were identified in the *NtJAZ1* and *NtJAZ3* promoters; **B,** electrophoretic mobility shift assay (EMSA) showed that PAP1 and TT8 bound to three MRE types and G-box of the NtJAZ1 and NtJAZ3 promoters, respectively. Competitive and non-competitive probes were used for binding of both TFs. 20× and 10×: concentrations of tested competitive and non-competitive probes 20 and 10 times as those MRE and G-box probes. **C,** fold change values from ChIP- qPCR showed that both PAP1 and TT8 bound to the *NtJAZ1* and *NtJAZ3* promoters *in vivo*. Approximately 150 bp *NtJAZ1* and *NtJAZ3* promoter fragments containing both MRE and G-box were enriched with anti-HA antibodies in qRT-PCR analysis. The regions of *NtJAZ1* and *NtJAZ3* promoters that do not contain a MRE and G-box were used as negative controls. Green and red bars represent wild type and red tobacco plants, respectively. **D,**schematic diagrams shows four effector constructs (PK2GW7- GFP, PAP1, TT8 and TTG1) and two reporter constructs (pGreenII-0800-Jaz1pro and Jaz3 pro) used for dual-luciferase assay. **E,** luciferase (LUC)/renilla (REN) luminescence ratios from dual luciferase assays showed that PAP1 and TT8 alone bound to and activated the promoters of both *NtJAZ1* and *NtJAZ3*, two TFs together increased promoter activity, and two TFs and WD40 together increased the highest activity. The promoters were fused to a firefly luciferase (reporter). The promoters of *NtJAZ1* and *NtJAZ3* genes were used in dual luciferase assay. Values represent the mean ± S.D. (n=5). Asterisks on top of bars indicate that the values are significantly higher than those of controls (*P<0.05, **P<0.01, ***P<0.001). Standard bars mean standard errors.

### Electrophoretic mobility shift assay of PAP1 and TT8 binding to promoters

To perform electrophoretic mobility shift assay (EMSA), we designed gene-specific primers (Supporting Table S1) for PCR to amplify the binding domains of both *PAP1* and *TT8*. The amplified *PAP1* binding domain from the N-terminus was 534 bp, which included the *R2R3 MYB* domain sequences for Gate-way cloning. The amplified *TT8* binding domain franking the *bHLH* region was 252 bp, in which both BamHI and SaclI restriction sites were added to two ends for further cloning. After the confirmation of sequence accuracy, three expression vector systems, pRSF-Dute, pet-Dute, and pDest-His-MBP were comparatively tested to induce soluble recombinant proteins. We selected the pDest-His-MBP vector, a Gateway cloning system, for the *R2R3* domain of *PAP1*. We found that the pRSF-Dute vector with a His-tag in the N-terminus was appropriate for expressing the bHLH domain of TT8 after several vectors were tested. As reported previously[69], detailed steps are listed in Method S1.

### Dual luciferase assays

Four reporter and four effector vectors were developed to perform dual luciferase assays that were completed to analyze the regulatory activity of PAP1 alone, TT8 alone, PAP1 and TT8 together, and PAP1/TT8/TTG1 (WD40) together in activating *NtJAZ1*, *NtJAZ3*, *PMT2,* and *ODC2* promoters. First, the pGreenII-0800 vector was used to develop four reporter vectors. Next, four effector vectors were developed from the PK2GW7 vector [20], to which the ORFs of *GFP*, *PAP1*, *TT8*, and *TTG1*were cloned. Each gene was driven by 35S promoter. Then, four reporter and four effector vectors were introduced into *Agrobacterium tumefaciens*strain GV3101, respectively. Last, the activated effector and reporter *Agrobacterium* GV3101 cells were used to inoculate leaves of *N. benthamiana* for transient expression assay. Detailed steps are described in Method S2.

### Cross-linked chromatin immunoprecipitation (ChIP) assays

A gateway cloning was used to develop two plasmids, in which *GFP* was fused to the 3’-end of *PAP1* and *TT8* ORFs without their stop codon for genetic transformation. A pair of primers (Supporting Table S1) was designed to amplify the *PAP1* ORF without its stop codon. The resulting PCR product was purified as described above and then cloned into the pDonr221 vector via a BP reaction. The *PAP1*fragment was further cloned to the upstream of *GFP* in the destination vector pGWB5 via a LR reaction. This cloning step generated a recombinant vector pGWB5-PAP1-GFP, in which *PAP1-GFP* was driven by 35S promoter. The pGWB5-PAP1-GFP vector was introduced into the competent *E. coli*Top10 cells. Positive colonies were selected on LB medium containing 50 mg/l kanamycin. One colony was used for suspension culture on the same shaker as described above. Cultured *E. coli*cells were harvested to isolate the pGWB5-PAP1-GFP binary plasmid, which was further introduced into competent *Agrobacterium* GV301 cells. To develop a binary vector expressing *TT8*, a pair of primers (Supporting Table S1) was designed for PCR. The reverse primer was designed to include a HA-tag sequence. The resulting PCR product was cloned to the plasmid PK2GW7 as described above. Next, the resulting binary vector PK2GW7-TT8-HA-tag was introduced into competent *Agrobacterium* GV301 cells.

### Agrobacterium

GV301 cells harboring *PAP-GFP* or *TT8-HA-tag* cDNA were activated and then injected to leaves of 5-week old wild type NL and KY171 seedlings that were grown in the phytotron. After the injection, plants were allowed to grow 72 hrs. The infected leaves were then harvested to liquid nitrogen to isolate nuclei and ChIP experiments. Detailed steps are listed in Method S3.

### Synthesis of T-DNA, development of binary constructs, and genetic transformation of two dark tobacco varieties

*PAP1* and *TT8* of *A. thaliana* encode a R2R3-MYB and bHLH TF, respectively. A DNA cassette was designed and then synthesized to stack *PAP1* and *TT8* for their coupled overexpression. From 5’ to 3’ end, the cassette was composed of *attL1*, the *PAP1* cDNA (NP_176057), *NOS* (terminator), *35S* promoter, the *TT8* cDNA (CAC14865), and *attL2* (Fig. S7 A). The synthesized sequence was introduced to the entry vector pUC57 and then cloned to the destination binary vector pK2GW7 by attL × attR combination (LR) reaction with LR clonase. The resulting binary vector, namely PAP1-TT8-pK2GW7 (Fig. S7 B), was further introduced into the competent cells of *Agrobacterium tumefaciens*strain GV3101. A positive colony was selected for genetic transformation of tobacco. In addition, the pK2GW7 vector was used for genetic transformation as control. These constructs were transformed into two globally commercial dark tobacco varieties, NL and KY171, respectively. Detailed steps (Fig. S7 C) for transformation and selection of T0 transgenic plants followed our protocols reported previously [20]. Transgenic plants from NL and KY171 were labeled as P+T-NL and P+T-KY, respectively. More than 20 T0 transgenic plants were obtained for each variety and grown in the greenhouse to select T0 seeds, 12 of which were further selected on medium supplemented with antibiotics to screen antibiotic resistant T1 progeny. A 3:1 ratio of seed germination to seed death indicated one single copy of T-DNA that was in the genomic DNA of T0 plants. This ratio was observed in the T1 progeny of T0 plants. In this study, we focused on lines with one single copy of transgenes. A large number of seeds were obtained from the T1 progeny of each line. Seeds from these types of T1 progeny were screened on an agar-solidified MS medium supplemented with 50 mg/l kanamycin. If all seeds from a T1 plant could germinate T2 seedlings, it indicated that this T1 plant was a homozygous line. Accordingly, red homozygous T2 plants (Fig. S7 M) were obtained and then used for growth in the greenhouse and field trial. In addition, PCR and qRT-PCR were performed to demonstrate the presence of transgene in the genome of T0 and T2 transgenic plants and the expression of both *PAP1*and *TT8* in both P+T-NL and P+T-KY plants.

### DNA and RNA extractions

Genomic DNA was extracted using a DNeasy Plant Mini Kit (QIAGEN, Hilden, Germany). RNA was extracted using Trizol extraction reagents. Fresh plant samples were ground into fine powder in liquid nitrogen. Extraction steps were followed the manufacturers’ protocols. Detailed steps are listed in Method S4.

### Polymerase chain reaction and quantitative reverse transcript-polymerase chain reaction

All real-time quantitative RT-PCR (qRT-PCR) analyses were performed for wild- type, vector control transgenic, and red transgenic plants. SYBR-Green PCR Master- mix (Bio-Rad, Hercules, CA) was used for qRT-PCR experiments on a Step-One Real-time PCR System (Thermo Fisher, Waltham, MA). Detailed steps are listed in Method S5.

### Assays of anthocyanin and proanthocyanidin in seedlings

Fresh leaves and roots (100 mg) of wild type, P+T-NL1, P+T-KY1, and vector control seedlings grown hydroponically (Supporting Figure S9) were used to extract anthocyanins and proanthocyanidins (PAs). Methods were followed our anthocyanin [20] and proanthocyanidin assay protocols [70]. Detailed steps are listed in Method S6.

### Field trial of P+T-NL1 and P+T-KY1 T2 progeny and air curing

Homozygous T2 progeny of P+T-NL1 and P+T-KY1 were planted at the research station in Oxford, North Carolina in 2015. In addition, NL and KY171 plants were planted as controls. Prior to planting transgenic plants in the field, we applied for GMO permits from USDA-APHIS for this field trial. North Carolina State University was licensed to grow transgenic tobacco plants in the field. Every single step strictly followed the protocol requested in the license. Field planting also followed the protocol of commercial tobacco production in North Carolina. These steps included seedling growth in a contained greenhouse, selection and design of field, planting, phenotypic observation, field management, topping, harvest of leaves, air curing, and cleanup of plant residues from the field (Table S2 and Supporting Figures S15-S19). Detailed steps are listed in Method S7 (Table S2, Figs S15-S19).

### Field trial of red PAP1 tobacco plants and flue-curing

PAP1 tobacco is the progeny of a homologous red PAP1 Xanthi plant (#292) that is generated by the overexpression of *PAP1* [21]. The field trials for PAP1 tobacco were conducted in 2011 and 2012. Field design, seed germination, plantation, field management, and plant growth management were the same as described above for P+T-NL1 and P+T-KY1 genotypes. The main differences were that leaves were harvested at different dates and flue-cured in barns. Detailed steps are listed in Method S8 (Fig. S22 A-E).

### Determination of TSNAs in cured leaves of field-grown P+T-NL1 and P+T-KY1 tobacco plants with high performance liquid chromatography coupled with triple quadrupole tandem mass spectrometer

Accurate quantification of tobacco specific nitrosamines (TSNA) was completed with high performance liquid chromatography coupled with triple quadrupole tandem mass spectrometer (HPLC-QQQ-MS/MS). This analysis was performed using industrial protocols on an Agilent 1200 series HPLC System coupled with AB Sciex Triple quadrupole mass spectrometer (API 5000 with Electrospray) at RJ Reynolds. Four TSNAs, N-nitrosonornicotine (NNN), N-nitrosoanatabine (NAT), N-nitrosoanabasine (NAB), and nicotine-derived nitrosamine ketone (4(methylnitrosamino)-1-(3-pyridyl)- 1-butanone, NNK) in air-cured leaves, were quantified. One hundred milligrams of dry leaf powder were extracted in an aqueous ammonium acetate solution (100 mM aqueous ammonium acetate containing four deuterium analogs for TSNA) and filtered through disposable PVDF syringe filters into autosampler vials. The extract was separated in a Phenomenex Gemini C18 column with 3.0 µm particle 2.0 x150 mm.

The injection volume was 2.0-5.0 µl. TSNAs were detected by multiple reaction monitoring (MRM) of the precursor ion to a product ion specific for each compound. Quantification was achieved using an internal standard calibration comprised of ten points. A separate internal standard was used for each analyte by using a mixture of four stable isotope-labeled analytes. Results are reported in units of ng/g (ppb). Data were determined to be acceptable if the correlation coefficient for the calibration curve was greater than 0.99, standards were within 85% to 115% of expected concentration values, check solution values were within 75% to 125% of target, peak shape and resolution were acceptable (based on historical data), and values for the QC samples were within established control limits. The instrument parameters for MS, HPLC gradient, the quadrupole mass spectrometer parameters, reagents and standards are listed in in Supporting Tables S3-S6.

### Determination of nicotine and other tobacco alkaloids in cured leaves of field grown P+T-NL1 and P+T-KY1 tobacco plants by GC

Analysis of gas chromatograph (GC) equipped with a flame ionization detector (FID) was performed using a standard industry method to quantify nicotine and other tobacco alkaloids at RJ Reynolds. In this study, analysis of nicotine and other tobacco alkaloids was performed on an Agilent 6890 gas chromatograph (GC) equipped with a flame ionization detector (FID) and an Agilent 7683 automatic sampler. One hundred milligrams of air-cured dry leaf powder were alkalinized in 2 mM sodium hydroxide (NaOH). This solution was extracted with methyl-tertbutyl ether (MTBE) spiked with an internal standard using a wrist-action shaker. The mixture was allowed to separate to two layers. The resulting MTBE layer was transferred to an autosampler vial for analysis of nicotine, anabasine, nornicotine, and myosmine with GC-FID. Quantification was achieved using an internal standard calibration comprised of six points. Data were determined to be acceptable if the correlation coefficient of the calibration curve was greater than 0.998, the response factors (RF) were consistent, the percent relative standard deviation (%RSD) for RF was equal or less than 5%, the check solution values were within 10% of target, values for the quality control (QC) samples were within established control limits, and chromatograms had appropriate identification of peaks. The instrument parameters, the oven program, and reagents and conditions used in alkaloid analysis are listed in Supporting Table S7-S9. The results were reported as percentiles (%, g/g, dry weight).

### Determination of TSNAs and tobacco alkaloids in flue-cured leaves of PAP1 plants with HPLC-ESI-MS

HPLC-ESI-MS analysis was performed on a 2010EV LC/UV/ESI/MS instrument (Shimadzu, Japan). One gram of powdered samples was extracted in 20 ml of methanol in a flask placed on a shaker for one hour at room temperature. The samples were centrifuged twice at 4000 rpm and aliquots of the supernatant were transferred to a 2.0 ml glass vial. The samples were separated on an Eclipse XDB-C18 analytical column (250 mm × 4.6 mm, 5 µm, Agilent, Santa Clara, CA, USA). The mobile phase solvents were composed of 1% acetic acid in water (solvent A; HPLC-grade acetic acid and LC–MS grade water) and 100% LC–MS grade acetonitrile (solvent B). To separate metabolites, the following gradient solvent system was used with ratios of solvent A to B: 90:10 (0–5 min), 90:10–88:12 (5–10 min), 88:12–80:20 (10–20 min), 80:20–75:25 (20–30 min), 75:25–65:35 (30–35 min), 65:35–60:40 (35–40 min), 60:40–50:50 (40–55 min), and 50:50–10:90 (55–60 min). The column was then washed for 10 min with 10% solvent B. The flow rate and the injection volume were 0.4 ml/min and 20 µl, respectively. The total ion chromatograms of positive electrospray ionization were recorded from 0 to 60 min by mass spectrum detector and mass spectra were scanned and stored from m/z of 120–1,600 at a speed of 1,000 amu/s. Standards of NAB, NAT, NNAL, NNK, and NNN (Sigma, St-Louis, MO) were used to establish standard curves with a coefficient value at least 98%. The peak values were used to estimate the levels of nicotine, nornicotine, anabasine, and anatabine.

### Determination of nicotine and nornicotine in leaves and roots of seedlings with HPLC-qTOF-MS

Seedlings of P+T-NL1, P+T-KY1, vector control, and wild type plants were hydroponically grown in the phytotron. Leaves and roots of 45-day old seedlings (Supporting Figure S9) were used to analyze nicotine and nornicotine in both leaves and roots. Fresh tissues were ground into fine powder in liquid nitrogen and dried in a 45°C oven. One hundred mg of powdered samples was extracted with 2.0 ml methanol. Other steps were as described above. The analysis of nicotine and nornicotine was performed with HPLC-qTOF-MS on Agilent 6520 time-of-flight LC-MS/MS instrument (Agilent Technologies, Santa Clara, CA, USA). Mobile solvents and column used for separation of metabolites were solvent A and B described above. The gradient solvent system was composed of ratios of solvent A to B: 95:5 (0–5 min), 95:5 to 90:10 (5–10 min), 90:10 to 85:15 (10–15 min), 85:15 to 45:55 (15–20 min), 45:55 to 25:75 (20–25 min), 25:75 to 95:5 (25–26 min). After the last gradient step, the column was equilibrated and washed for 10 min with solvents A: B (95:5). The flow rate was 0.4 ml/min. The injection volume of samples was 5.0 µl. The dry gas flow and the nebulizer pressure were set at 12 l/min and at 50 psi, respectively. Metabolites were ionized with the negative mode. The mass spectra were scanned from 100 to 3000 m/z. The acquisition rate was three spectra per second. Other MS conditions included fragmentor: 150 V, skimmer: 65 V, OCT 1 RF Vpp: 750 V, and collision energy: 30 psi. Standard curves were developed for both nicotine and nornicotine. The results were reported as mg/g, dry weight.

### Statistical analysis

Metabolite values for each group of samples such as B1 were averaged from samples of three plots. One-way ANOVA was used to evaluate the statistical significance of the contents of nicotine, nornicotine, anabasine, NNN, NNK, NAT, and NAB between red and wild type leaves. Standard deviation was used to show the variation of metabolite contents. For gene expression, protein analysis, nicotine and nornicotine contents in seedlings, all values are averaged from four or five biological replicates. Student t-test was performed to evaluate statistical significance. P-values less than 0.05, 0.01, and 0.001 mean significant, very significant, and extremely significant.

## Results

### MRE and G-box elements identified in the promoters of NtJAZ1, 3, 7, and 10

NtJAZ1 and NtJAZ3 are two main repressors of nicotine biosynthesis [54, 55, 60]. In recent, totally 15 *NtJAZs* were mined from the tobacco genomes, showing that *NtJAZ7*, and *NtJAZ10* were also associated with the regulation of nicotine biosynthesis [61]. We identified the promoter sequences of 13 *NtJAZs* from the genome of NC90 tobacco curated at NCBI and named these as *NtJAZ1pro* through *NtJAZ13pro*. One or 1.5 kb nucleotides in the proximal upstream from the start codon (ATG) were obtained for sequence characterization. Sequence analysis revealed that *NtJAZ1pro*, *NtJAZ3pro*, and *NtJAZ7pro* had both MYB recognizing element (MRE) and G-box elements, while *NtJAZ10pro* only had G-box elements (Figs S1-S2). Next, we cloned the promoter sequences of *NtJAZ1pro* and *NtJAZ3pro* from two commercial smokeless dark tobacco cultivars, <u>N</u>arrow <u>L</u>eaf Madole (NL) and <u>KY</u>171 (KY). Their sequences were identical to those of *NtJAZ1pro* and *NtJAZ3pro* from TN90 (Fig. S1), therefore, we used the same names in this report. Sequence analysis further substantiated the MRE and G-box elements in *NtJAZ1pro*and *NtJAZ3pro*.

*NtJAZ1pro* has two MRE and two G-box elements (Fig. 2 A and Fig. S1). The two MRE elements are MRE1: <u>ACCCACC</u> at position -180 and MRE2: <u>ACCCCAC</u> at position -172. The two G-boxes are <u>CACGTG</u> at positions -232 and AACGTG at -395. *NtJAZ3pro* has one MRE3 element: <u>AACTACC</u> at position -169 and one G-box: <u>CACGTG</u> at position -960 (Fig. 2 A and Fig. S1).

### EMSA shows the binding of PAP1 and TT8 to the NtJAZ1 and NtJAZ3 promoters in vitro

Electrophoretic mobility shift assay (EMSA) was carried out to test whether PAP1 and TT8 could bind to the MRE and G-box elements of *NtJAZ1pro and NtJAZ3pro in vitro*. The R2R3-MYB binding domain (amino acids 1-178) of Arabidopsis PAP1 was cloned to the pDEST-HisMBP vector, in which it was fused with the His-MBP-Tag at the N-terminus (Fig. S3 A). The bHLH binding domain (amino acid 359-443) of TT8 was cloned to the PRSF-Dute vector, in which it was fused with the His-Tag at the N-terminus (Fig. S3 B). Two constructs were introduced to *E. coli* to induce recombinant proteins, which were further purified with a His-tag purification system (Fig. S3 B). Biotin probes were prepared for MRE1 (ACCCACC), MRE2 (ACCCCAC), MRE3 (AACTACC), and G-box (CACGTG). Both competitive and non-competitive probes were used as controls for EMSA. The resulting data showed that the recombinant R2R3 binding domain directly bound to three MRE element probes and the recombinant bHLH binding domain directly bound to the G-box elements (Fig. 2 B). The binding activity of both recombinant R2R3 and bHLH binding domains was competed by competitive probes but not competed by non-competitive probes (Fig. 2 B). These data demonstrate that both PAP1 and TT8 bind to the *NtJAZ1pro and NtJAZ3pro*.

### ChIP-Q-PCR shows the binding of PAP1 and TT8 to the NtJAZ1 and NtJAZ3 promoters in vivo

ChIP-Q-PCR analysis was performed to examine whether PAP1 and TT8 could bind to *NtJAZ1pro* and *NtJAZ3pro in vivo*. In ChIP experiments, *PAP1* was fused at the 5’- end of *GFP* in the binary vector pGWB5 to obtain pGWB5-PAP1-GFP, which was introduced into leaves of wild type NL and KY171 seedlings for transient expression via injection of *Agrobacterium*. *TT8* was fused at the 5’-end of an *HA-tag* sequence in the binary vector PK2GW7 to obtain PK2GW7-TT8-HA-tag, which was also introduced into leaves of wild type NL and KY171 seedlings for transient expression by injection of *Agrobacterium*. The IP experiments of PAP1 and TT8 were performed by using GFP and HA monoclonal antibodies. The ChIP PCR results showed that the amplified *NtJAZ1* and *NtJAZ3* promoter fragments were increased 5.0-40.4 times by PAP1 and 6.1.8-37.5 times by TT8 (Fig. 2 C).

### Dual-luciferase assay demonstrates the activation of NtJAZ1 and NtJAZ3 promoters by PAP1, TT8, PAP1/TT8, and PAP1/TT8/TTG1

Dual-luciferase assay was completed to demonstrate whether PAP1 alone, TT8 alone, PAP1 and TT8 together, and PAP1/TT8/TTG1 (WD40) together could activate the *NtJAZ1* and *NtJAZ3* promoters in tobacco. PAP1, TT8, and TTG1 were used as effectors. The ORF sequences of *PAP1*, *TT8*, *TTG1*, and *GFP* were respectively cloned into the PCB2004 plasmid to generate four constructs, in which each gene was driven by a 35S-promoter (Fig. 2 D). The *NtJAZ1pro* and *NtJAZ3pro* were cloned into the plasmid pGreenII-0800 to drive the luciferase gene to obtain *NtJAZ1pro-*luciferase and *NtJAZ3pro*-luciferase reporters (Fig. 2 D). Effectors and reporters were incubated for dual-luciferase assays. Promoters alone and a binary vector were used as controls. The resulting data showed that compared with controls, PAP1 and TT8 alone enhanced the activity of *NtJAZ1pro* 3.8 and 7.2 times, respectively, and increased the activity of *NtJAZ3pro* 1.9 (slightly) and 2.1 (significantly) times, respectively (Fig. 2 E). PAP1 and TT8 together enhanced the activity of *NtJAZ1pro* and *NtJAZ3pro* approximately 4.0 and 6.3 times (Fig. 2 E). Furthermore, PAP1, TT8, and TTG1 together increased 12.7 and 15.6 times of the activity of both *NtJAZ1pro* and *NtJAZ3pro*, much higher than PAP1 alone, TT8 alone, and PAP1 and TT8 together (Fig. 2 E). These results demonstrate that PAP1 and TT8 alone, PAP1 and TT8 together, PAP1-TT8-TTG1 complex transcriptionally activate both *NtJAZ1* and *NtJAZ3*.

### PAP1, TT8, PAP1/TT8, and PAP1/TT8/TTG1 do not activate promoters of ODC2 and PMT of tobacco

*NtODC2* and *NtPMT2* are two key upstream pathway genes of the nicotine biosynthesis. We obtained 1.0 kb long sequences for these two genes’ promoters from the tobacco genome database curated at NCBI. The promoter sequence of *NtPMT2* has an MRE-like element (<u>AACAACC</u> at position -530) and a G-box like (<u>CACGTT</u> at position -186) element, while the sequences of the *NtODC2*promoter does not have MRE and G-box elements (Fig. S4).

EMSA experiments were performed to test whether PAP1 and TT8 could bind to MRE-like and G-box-like elements. The EMSA results indicated that the TT8 binding domain could bind to the G-box-like (CACGTT) element. However, when a competitive probe was added, the binding signal was lost (Fig. S5 A). Furthermore, compared with the positive control G-box elements from *NtJAZ1* and *NtJAZ3* promoters, the resulting data showed that the signal from G-box like element was much weaker (Fig. S5 A). This result indicates that TT8 can weakly bind to the promoter of *NtPMT2*. When the MRE-like element probe was prepared for EMSA, the resulting data showed that PAP1 could not bind to the MRE-like probe (<u>AACAACC</u>), while PAP1 strongly bound to the positive control MRE1 and MRE3 probes of *NtJAZ1* and *NtJAZ3* promoters (Fig. S5 B).

Dual-luciferase assay and ChIP-Q-PCR were performed to test whether PAP1 and TT8 could activate the *ODC2* and *PMT2* promoters. The dual-luciferase assay data did not show that PAP1 alone, TT8 alone, PAP1/TT8 together, and the PAP1/TT8/TTG1 complex could activate the activity of the two promoters (Fig. S6 A). Meanwhile, the ChIP-Q-PCR data showed that PAP1 and TT8 could not enrich binding elements of the two promoters *in vivo* (Fig. S6 B). These data indicate that PAP1, TT8, PAP1/TT8, and PAP1/TT8/TTG1 cannot activate the promoters of these two key pathway genes.

### Engineering of red genotypic P+T-NL and P+T-KY tobacco plants

An expression cassette was synthesized to stack the ORF sequences of *PAP1* (P) and *TT8*(T) and then used to generate 20 T0 transgenic plants from each of Narrow Leaf Madole (NL) and KY171 (KY), two major commercial varieties (Fig. S7 A, B, C, E, and G-H). In addition, vector control transgenic plants were generated (Fig. S7 D and F). The first 12 red T0 transgenic plants from the two cultivars were tagged as P+T- NL1 to 12 and P+T-KY1 to 12, respectively. The first 12 T0 vector control transgenic plants from the two cultivars were tagged as NL1 to 12 and KY1 to 12, respectively. These T0 plants were selected to obtain seeds to screen T1 and homozygous T2 progeny (Fig. S7 I-M), during which the copy number of the transgenes was determined. Five P+T-NL lines (1-5) and five P+T-KY lines (1-5) were confirmed to contain one single copy of the stacked two transgenes. Quantitative RT-PCR showed that homozygous T2 progeny highly expressed *PAP1* and *TT8* (Fig. S8). Furthermore, qRT-PCR was performed to understand the expression of *CHS*, *CHI, F3H, F3’H*, *DFR*, and *ANS* involved in the biosynthesis of both anthocyanin and proanthocyanidin (PA) and the expression of *3-GT* involved in the glycosylation of anthocyanidins (Fig. 1 A) in T2 seedlings (Fig. S9). The resulting data showed that the expression levels of these seven genes were up-regulated 2-1000 times (Fig. S10 AB) in pink roots and red leaves of the P+T-NL and P+T-KY genotypes. Further UV spectrometry-based measurements showed that the roots and leaves of analyzed P+TNL1 and P+T-KY1 genotypes produced anthocyanins, while those of control plants did not (Fig. S11). It was noticeable that the levels of anthocyanins were significantly higher in leaves than in roots of both P+T-NL1 and P+T-KY1 plants (Fig. S11). To understand this differentiation, the expression of *ANR* specifically involved in the colorless PA formation [71] was analyzed. The resulting data showed that the expression of *ANR* was significantly activated in roots but not in leaves of the two red genotypes (Fig. S10). To substantiate this, root extracts of both P+T-NL1 and P+T-KY1 seedlings were treated with DMACA. A bluish color was immediately developed from the reaction (Fig. S12 A-B), showing the presence of PAs. However, this bluish color was not developed from the extracts of both P+T-NL1 and P+T-KY1 leaves and wild type roots and leaves, indicating the lack of PAs. Further measurement of DMACA-treated samples at 640 nm showed that the ABS values were 5-6 times higher in the root extracts than in the leaf extracts of P+T-NL1 and P+T-KY1 and in the leaf and root extracts of wild type (Fig. S12 C-D). These data showed that the less red pigmentation in roots resulted from the conversion of red anthocyanidins to colorless PAs by ANR. In addition, the less red pigmentation in roots might result from the lack of light in the hydroponic condition and secretion of anthocyanins.

### Upregulation of NtJAZ1, 3, 7, and 10 and downregulation of the nicotine biosynthesis in red P+T-NL and P+T-KY plants

Both transcriptional and metabolic profiling were performed to characterize the biosynthesis of nicotine in homozygous T2 seedlings (Fig. S9) of different genotypes selected, including P+T-NL1, P+T-NL2, P+T-NL3, P+T-KY1, P+T- KY 2, and P+T- KY3. Quantitative RT-PCR was performed to characterize the transcriptional profiles of 16 genes (Fig. 1 B) involved in the nicotine biosynthesis. The expression levels of *NtJAZ1, NtJAZ3*, *NtJAZ7,* and *NtJAZ10* were significantly increased in roots of both P+T-NL and P+T-KY genotypes (Fig. 3 A-B and K-L, Fig. S13). The expression levels of *PMT1*, *PMT2*, *MPO*, *QPT2*, *A622*, *BBL* and *ERF189* were significantly reduced in roots of both P+T-NL (Fig. 3 D-J) and P+T-KY plants (Fig. 3 N-T). The expression levels of *OCD2* was not altered in the P+T-NL1, 2, and 3 lines (Fig. 3 C) but significantly decreased in the P+T-KY1, 2 and 3 lines (Fig. 3 M), indicating the different responses of this gene expression in two commercial varieties to transgenes. The expression levels of *MYC1a*, *MYC1b*, *MYC2a*, and *MYC2b* were not altered in roots of both P+T-NL and P+T-KY plants (Fig. S14). HPLC-MS analysis showed that the contents of nicotine, nornicotine, and the total amount of the two compounds in roots and leaves of P+K-NL1, P+K-NL2, P+K-NL3, P+K-KY1, P+K-KY2, and P+K- KY3 plants were significantly reduced 1.5-3.4, 1.3-2.5, and 1.9-3.3 fold, respectively (Fig. 4).

**Figure 3.**
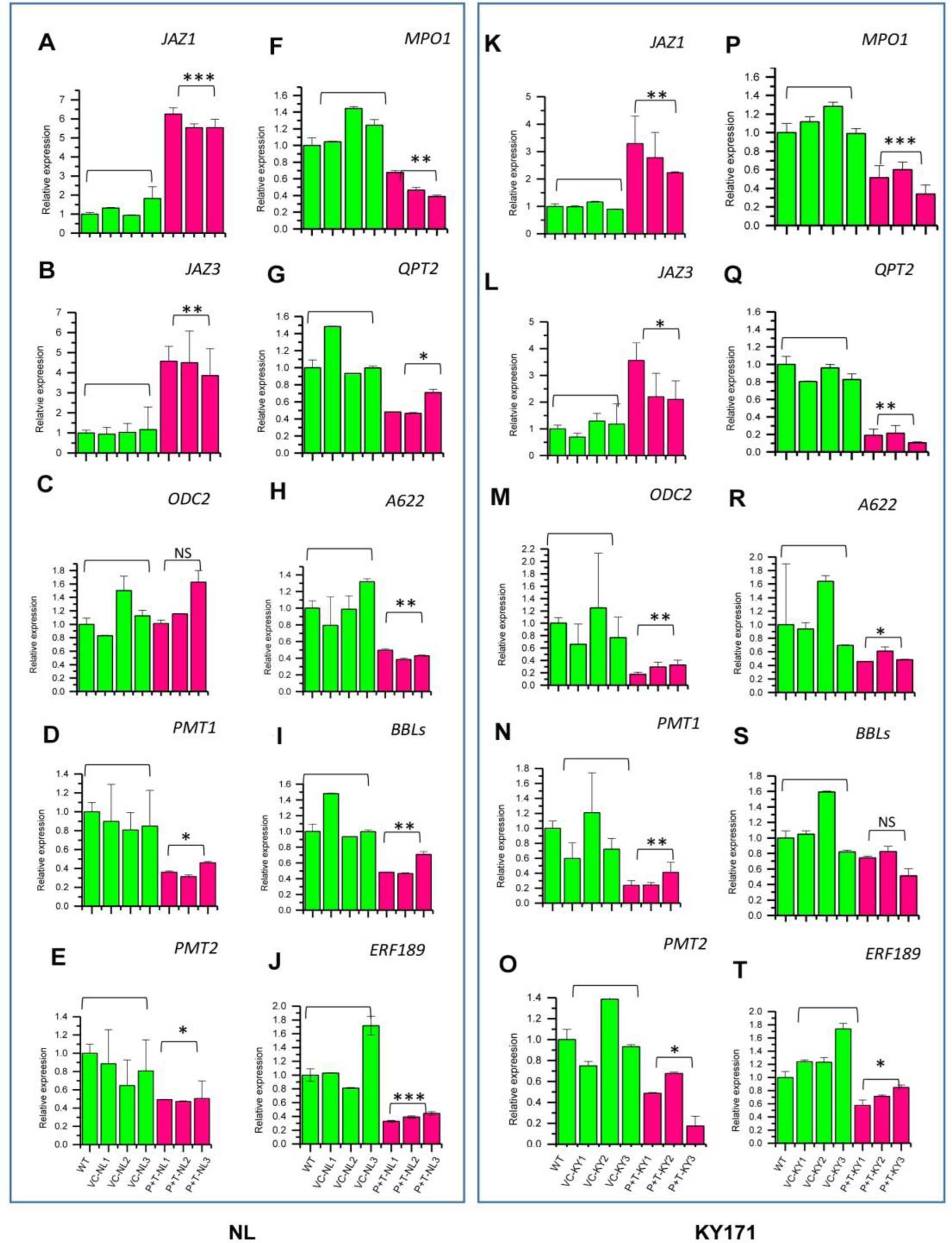
Upregulation of two *NtJAZs* and downregulation of nicotine pathway genes and *ERF189* in roots of P+T transgenic seedlings. **A-J**, in roots of P+T-NL transgenic lines, the expression levels of *NtJAZ1* (**A**) and *NtJAZ3* (**B**) were significantly upregulated; the expression level of *ODC2* (**C**) was not changed; while the expression levels of *PMT1* (**D**), *PMT2* (**E**) *MPO* (**F**), *QPT* (**G**), *A622* (**H**), *BBLs* (**I**) (primer pairs designed for all three BBLs), and *ERF189* (**J**) were significantly downregulated. **K-T**, in roots of P+T-KY transgenic lines, the expression levels of *NtJAZ1* (**K**) and *NtJAZ3* (**L**) were significantly upregulated, while, the expression levels *ODC2* (**M**), *PMT1* (**N**), *PMT2* (**O**) *MPO* (**P**), *QPT* (**Q**), *A622* (**R**), *BBLs* (**S**), and *ERF189* (**T**) were significantly downregulated. Data of three transgenic lines are shown for both P+T-NL and P+T-KY genotypes. Wild type samples were pooled from five plants. Data of three lines are shown for vector control transgenic NL and KY171 plants. WT: wild type, P+T NL1, P+T NL2, P+T NL3: three lines of stacked *PAP1* and *TT8* transgenic NL plants, VC-1, 2, and 3-NL: three vector control transgenic NL lines, P+T KY1, P+T KY2, P+T KY3: three lines of stacked *PAP1*and *TT8* transgenic KY171 plants. VC-1, 2, and 3-KY: three vector control transgenic KY171 lines. Green and red bars represent wild type and vector control transgenic, and red P+T-NL and P+T-KY plants, respectively. Values represent the mean ± S.D. (n=3). Asterisks on top of the bars indicate that values are significantly lower or higher in red transgenic lines than in wild type plants (*P<0.05, **P<0.01, ***P<0.001). Standard bars mean standard errors.

**Figure 4.**
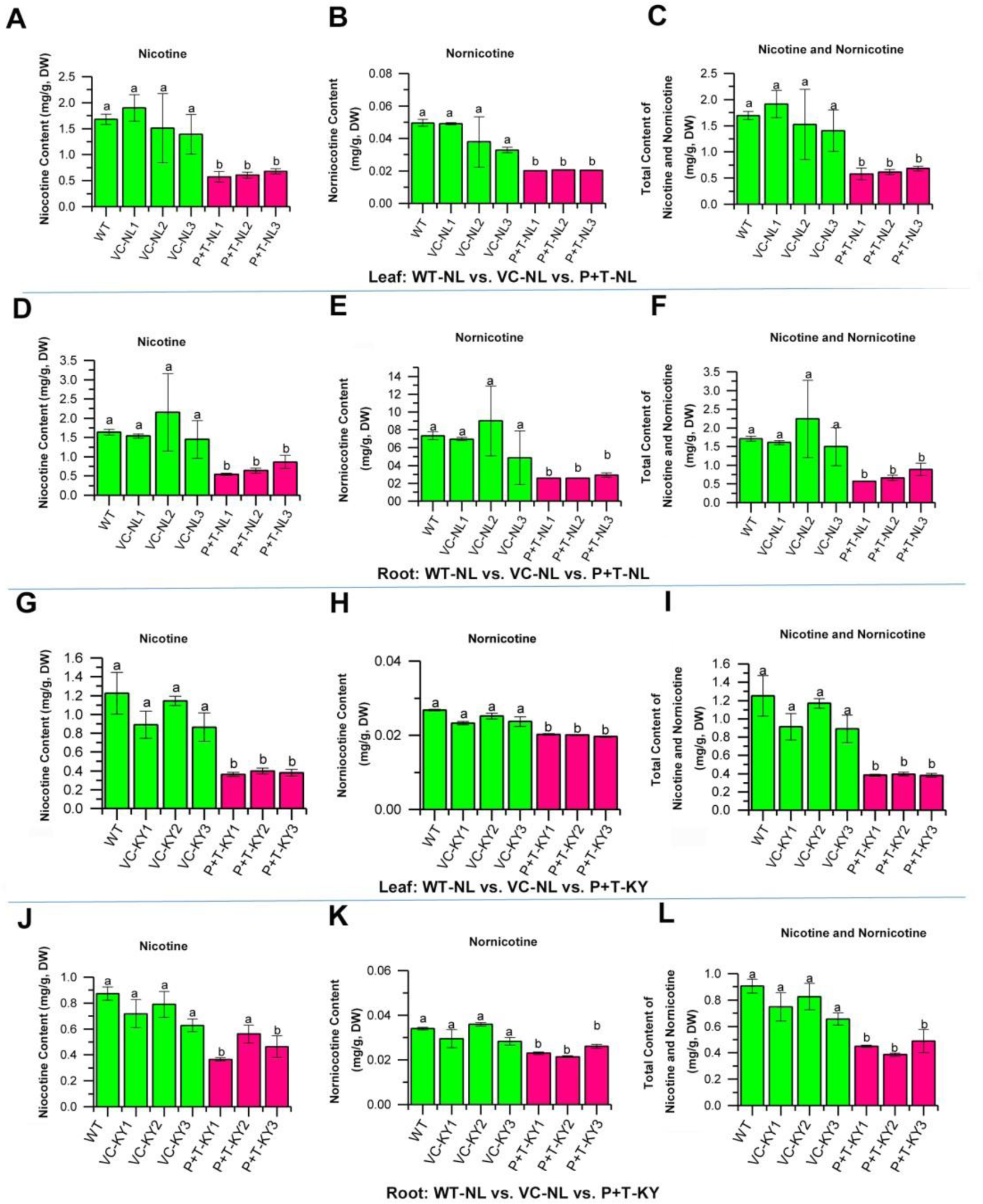
Significant reduction of nicotine and nornicotine contents in leaves and roots of P+T-NL and P+T-KY seedlings. The contents of nicotine and nornicotine were significantly reduced in roots and leaves of both transgenic P+T-NL and P+TKY lines. **A-C:** reduction of nicotine (**A**), nornicotine (**B**), and total nicotine and nornicotine (**C**) in leaves of three P+T-NL lines (1, 2, and 3); **D-F:**reduction of nicotine (**D**), nornicotine (**E**), and total nicotine and nornicotine (**F**) in roots of three P+T-NL lines (1, 2, and 3); **G-I:** reduction of nicotine (**G**), nornicotine (**H**), and total nicotine and nornicotine (**I**) in leaves of three P+T-KY lines (1, 2 and 3); **J-L:** reduction of nicotine (**J**), nornicotine (**K**), and total nicotine and nornicotine (**L**) in roots of three P+T-NL lines (1, 2, and 3). Bars labeled with “a” and “b” means p- value less than 0.05 and bars labelled with the same lowcase “a” or “b” means no significant difference.

### Field trials and reduction of nicotine, nornicotine, anatabine, anabasine, myosmine, and four TSNAs in red P+T-NL1 and P+T-KY1 tobacco plants

Field trials that follow the commercial production practices of tobacco are essential to examine the effects of new varieties or genes on the contents of nicotine, other tobacco alkaloids, and TSNAs. In addition, field farming practices were necessary to test the agricultural significance of this *de novo* regulation design. The field practice exactly followed the protocols of both industry tobacco and GMO tobacco growth.

We planted T2 homozygous P+T-NL1 and P+T-KY1 progeny and their corresponding wild-type tobacco variety in the field at the research station in Oxford, North Carolina (Fig. 5 A-C and Figs S15-S18). Tobacco leaves were air-cured in a contained ventilation barn (Fig. 5 D, Figs S18-S19).

**Figure 5.**
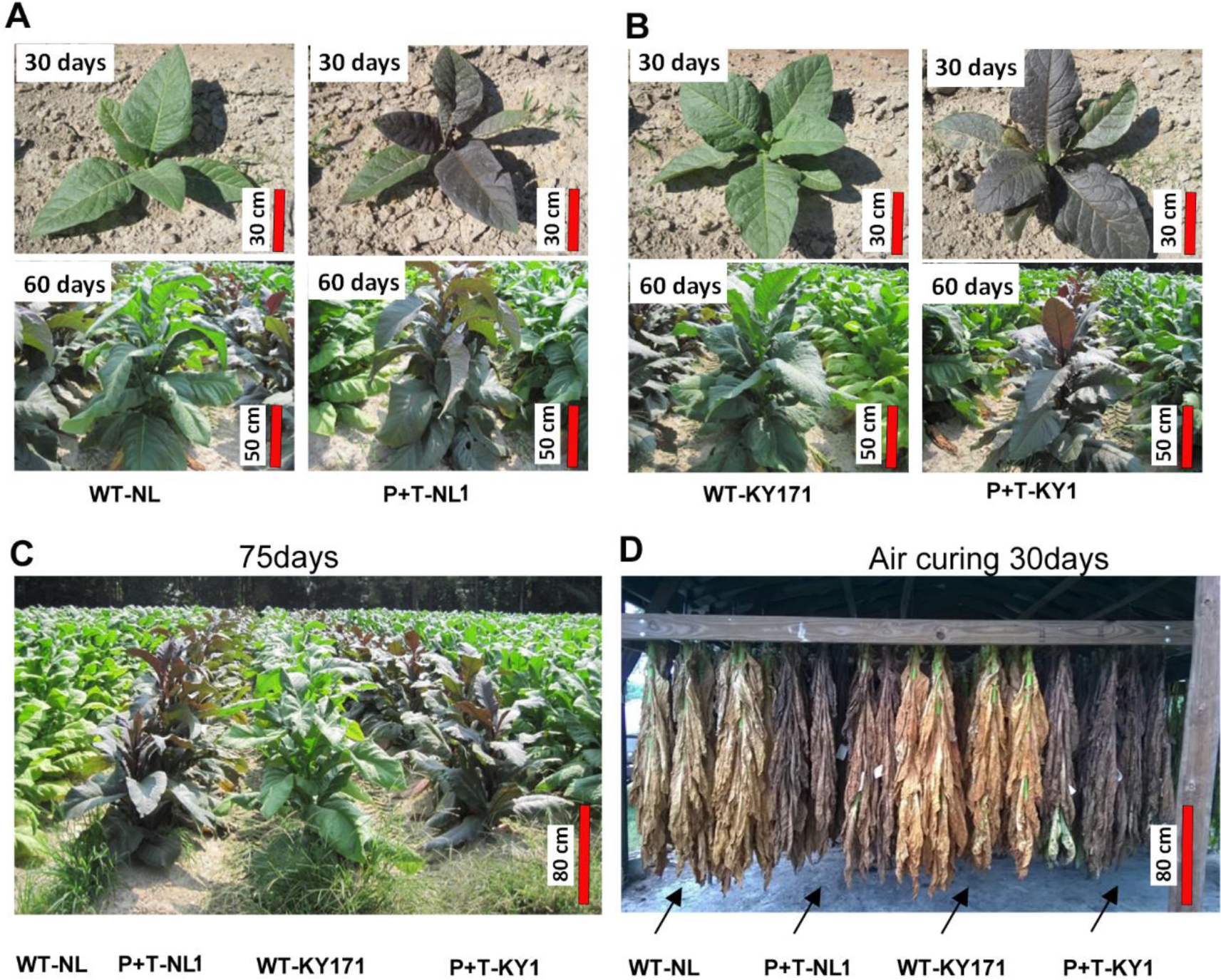
Phenotypes of P+T-NL1 and P+T-KY1 plants versus their corresponding wild-type tobacco plants in the field and of air-cured leaves. Field farming practice of four genotypes was performed in Oxford, North Carolina. **A**, phenotypes of 30-day and 60-day old WT-NL vs. red P+T-NL1 plants. **B**, phenotypes of 30-day and 60-day old WT-KY171 vs. red P+T-KY1 plants. **C-D**, phenotypes of topped plants (**C**) and leaves in air curing (**D**). Plant name abbreviations are WT-NL: wild-type Narrow Leaf Madole (NL) variety, P+T-NL1: Stacked *PAP1* and *TT8* transgenic NL line 1, WT-KY171: wild-type KY171 variety, and P+T-KY1: stacked *PAP1* and *TT8* transgenic KY line 1.

Quantification with GC-FID showed the significant reduction of nicotine, nornicotine, anabasine, anatabine, and myosmine in most or all cured leaves of red P+T-NL1 and P+T-KY1 tobacco plants compared to those of non-transgenic controls. The contents of nicotine were significantly reduced by 45-51% in all cured leaf groups of P+T-NL1 (Fig. 6 A) and by 19-30% in upper three groups of cured leaves of P+T-KY1 (Fig. 6 B). The contents of nornicotine were significantly decreased by 39-44% and 30-40% in all cured leaf groups of T-P-NL1 and P+T-KY1, respectively (Fig. 6 C and D). The contents of anabasine, anatabine, and myosmine were significantly reduced by 27- 45%, 55-66%, and 19-25% in all cured leaf groups of P+T-NL1 (Fig. 6 E, G, and I), respectively, and were significantly reduced by 27-45%, 3256%, and 29-35% in all cured leaf groups of P+T-KY1, respectively (Fig. 6 F, H, and J).

**Figure 6.**
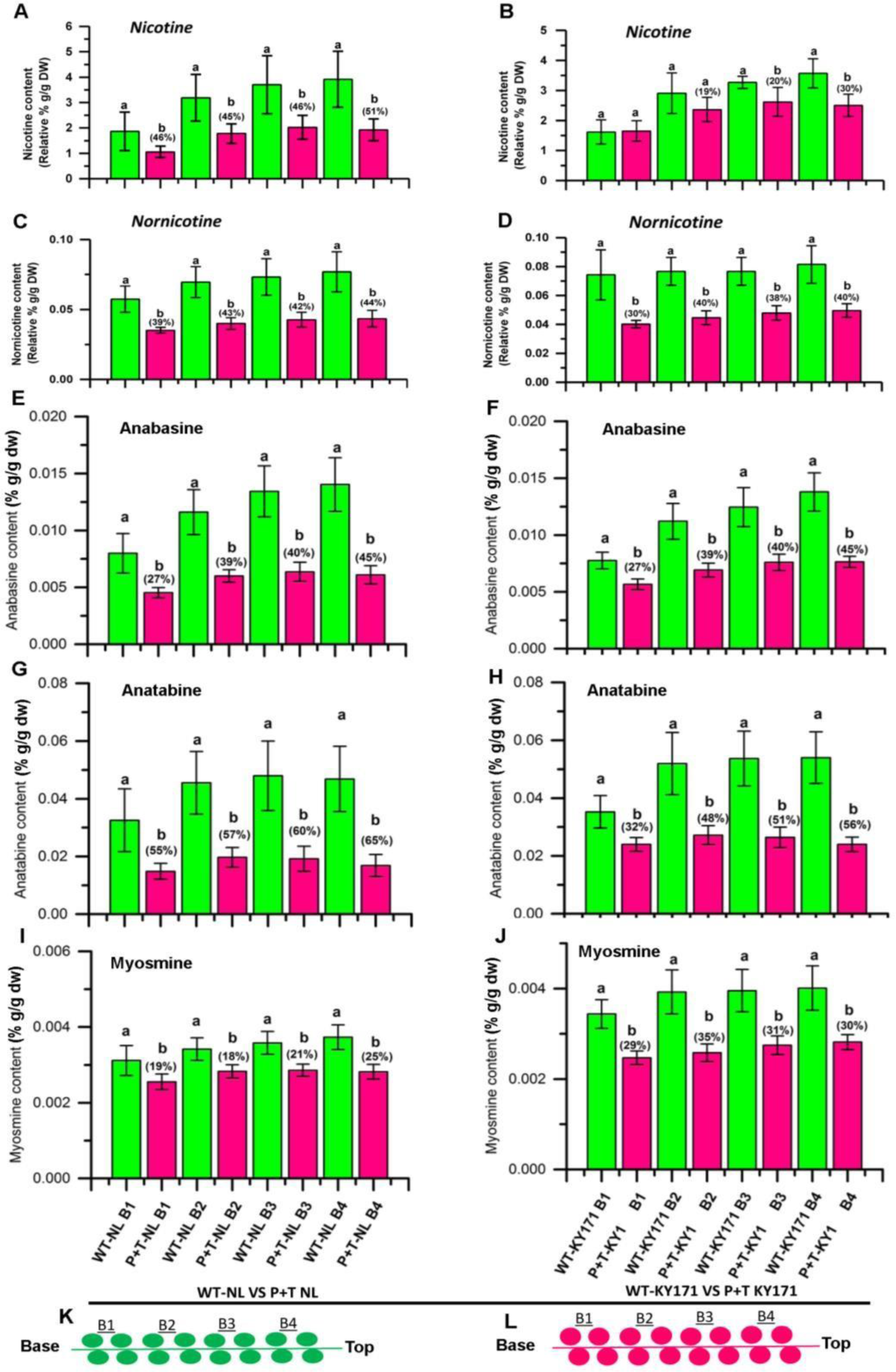
Reduction of nicotine and four other alkaloids in cured leaves of red P+T-NL1 and P+T-KY1 tobacco plants. **A-J**, reduction of nicotine (**A-B**), nornicotine (**C-D**), anabasine (**E-F**), anatabine (**G-H**), and myosmine (**I-J**) in four leaf groups of P+T-NL (**A, C, E, G, and I**) and P+T-KY plants (**B, D, F, H, and J**). **K-L**, cartoons showing leaf groups harvested from both wild type (**K**) and transgenic red plants (**L**). Green and red bars represent wild type and red tobacco plants, respectively. B1, B2, B3, and B4 labels represent the 1^st^, 2^nd^, 3^rd^, and 4^th^ group of leaves from the base to the top of tobacco plant, respectively. Each metabolite in each leaf group was quantified from leaves harvested from 120 tobacco plants in three plots. One-way ANOVA test included in the SAS software was used to evaluate statistical significance. Bars labelled with lowcase “a” and “b” mean significant difference between paired green and red bars (n=120, P-value less than 0.05). Percentages labeled in the top of red bars mean reduction compared to green bars. The standard bars mean standard deviation. Abbreviations, WT-NL: wild-type Narrow Leaf Madole (NL) variety, P+T-NL1: Stacked *PAP1* and *TT8* transgenic NL line 1, WTKY: wild-type KY171 variety, and P+T-KY1: stacked *PAP1* and *TT8* transgenic KY line 1.

Quantification with HPLC-QQQ-MS/MS showed that the contents of four carcinogenic TSNAs were significantly diminished in most or all leaf groups of P+TNL1 and P+T-KY1 plants. The content of NNN was significantly decreased by 63-74% to a level of 0.136-0.375 ppm in all four groups of cured P+T-NL1 leaves and by 67-79% to a level of 0.208-0.552 ppm in three groups of cured P+T-KY1 leaves (Fig. 7 A and B). The content of NNN in the B3 leaf group of P+T-KY1 was reduced by 63% to 0.488 ppm with a slight significance. The content of NNK was reduced by 38-72% to a level of 0.1-0.173 ppm and by 30-74% to a level of 0.151-0.3 ppm in four groups of cured P+T-NL1 and P+T-KY1 leaves, respectively, in which the reduction in three leaf groups of each genotype was significant (Fig. 7 C and D). The content of NAT was significantly decreased by 77-92% to a level of 0.093-0.159 ppm and by 70-80% to a level of 0.159-0.285 ppm in all leaf groups of P+T-NL1 and P+T- KY1, respectively (Fig. 7 E and F). The content of NAB was significantly decreased by 72-80% to a level of 0.011-0.021 ppm and by 65-77% to a level of 0.015-0.031 ppm in all leaf groups of P+T-NL1 and P+T-KY1, respectively (Fig. 7 G and H).

**Figure 7.**
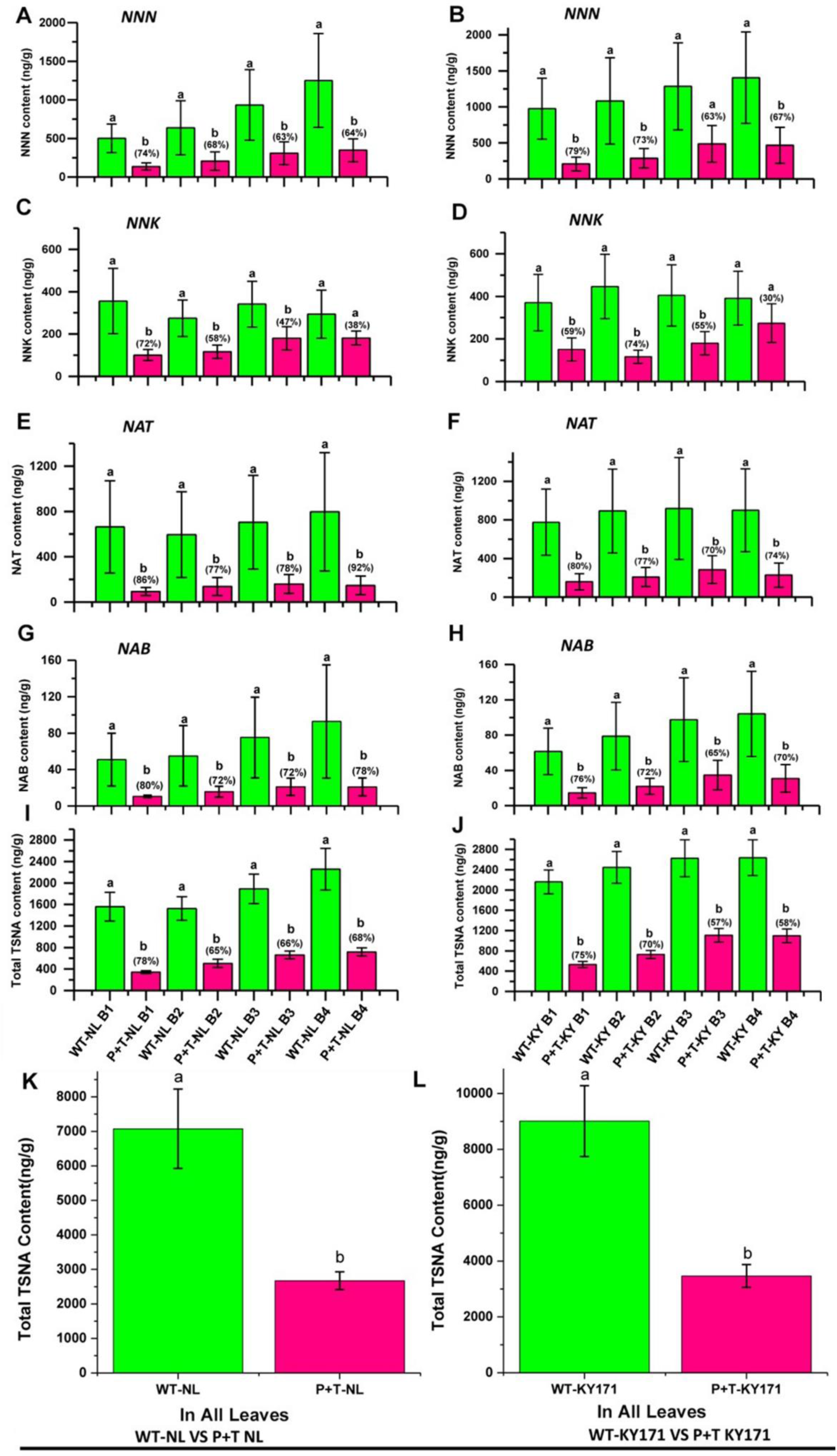
Reduction of four individual and total TSNAs in leaves of red P+T-NL1 and P+T-KY1 tobacco plants. **A-H,** reduction of NNN (**A-B**), NNK (**C-D**), NAT (**E-F**), and NAB (**G-H**) in each leaf group of P+T-NL1 (**A, C, E,** and **G**) and P+TKY1 (**B, D, F,** and **H**) plants; **I-J,** reduction of total TSNA contents summed from four individual TSNAs of each leaf group of P+T-NL1 (**I**) and P+T-KY1 (**J**) plants; **K-L,** reduction of total TSNA contents summed from four individual TSNAs of all leaves of P+T-NL1 (**K**) and P+T-KY1 (**L**) plants. Green and red bars represent wild type and red tobacco plants, respectively. B1, B2, B3, and B4 labels represent the 1^st^, 2^nd^, 3^rd^, and 4^th^ group of leaves from the base to the top of plants, respectively. Each metabolite in each group of leaves was quantified from 120 tobacco plants. One-way ANOVA test included in the SAS software was used to evaluate statistical significance. Paired green and red bars labelled with lowcase “a” and “b” mean significant difference (n=120, P-value less than 0.05). Percentages labeled on the top of red bars indicate reduction levels compared to green bars. Abbreviation, WT-NL: Narrow Leaf Madole, WT-KY171: wild type KY171, P+T-NL: stacked PAP1 and TT8 transgenic NL line 1, and P+T-KY1: stacked PAP1 and TT8 transgenic KY171 line 1.

Furthermore, the contents of NNN, NNK, NAT, and NAB were summed to obtain the total TSNA contents in each leaf group. The total contents of four TSNAs in each group of wild type NL and KY171 leaves were 1.6-2.2 ppm and 2.2-2.6 ppm, while these values were significantly reduced to be 0.4-0.6 ppm and 0.6-1.0 ppm in each group of P+T-NL1 and P+T-KY1 leaves (Fig. 7 I-J). The average contents of four TSNAs in the entire plant leaves of wild type NL and KY171 were approximately

1.77 ppm and 2.25 ppm, while these values were significantly reduced to be 0.69 ppm and 0.87 ppm in those of P+T-NL1 and P+T-KY1 (Fig. 7 K-L).

Furthermore, the total contents of tobacco alkaloids in each group of leaves were summed from nicotine, four other alkaloids, and total TSNAs. The resulting data showed that the contents of total alkaloids were significantly decreased by 44-51% in four leaf groups of P+TNL1 and 22-35% in the upper three group leaves of P+T-KY1 (Fig. S20 A-B).

### Field trials and reduction of nicotine, nornicotine, anatabine, anabasine, myosmine, and four TSNAs in flue-cured leaves of the red PAP1 tobacco

Our red PAP1 tobacco genotype (Fig. S22 B) is a homozygous red variety generated from Xanthi by the overexpression of *PAP1* [21]. Gene expression analysis showed that the expression of *PAP1*upregulated *NtAn1a* and *NtAn1b* (Fig. S21), two *TT8* homologs [72]. This result indicated that *PAP1* alone activated tobacco anthocyanin biosynthesis via upregulating the endogenous *N. tabacum TT8*(*NtTT8*) homologs. Therefore, to substantiate our *de novo* design, we used this genotype to test whether PAP1 and activated NtTT8 homologs could reduce nicotine, other tobacco alkaloids, and TSNAs.

Two years’ field trials were completed. The field design (Fig. S22 A-B) and cropping management were the same as described above. When leaves were harvested, they were harvested into five groups (Fig. S22 C-D) and then flue-cured in barns (Fig. S22 E). Analysis with HPLC-ESI-MS obtained peak values that showed the reduction of nicotine, nornicotine, anabasine, and anatabine. The levels of nicotine were significantly or slightly reduced by 10-50% and 5-25% in each group of leaves in 2011 and in 2012, respectively (Fig. S23 A-B). The levels of nornicotine, anabasine, and anatabine were significantly or slightly reduced in all or most group leaves by 25- 70% and 5-55%, 30-60% and 5-20%, and 5-60% and 10-40% in 2011 and 2012, respectively (Fig. S23 A-B). In addition, the total cumulative peak level of each alkaloid was summed from all leaf groups of the entire PAP1 tobacco plant. The resulting data showed that the total cumulative levels of nicotine, nornicotine, anabasine, and anatabine in the entire plant were significantly reduced by 30%, 45%, 50%, and 40% in 2011 and 10%, 20%, 10%, and 30% in 2012 respectively (Fig. S24 A-B).

The contents of NNN, NNK, NAB, and NAT were reduced in flue-cured leaf groups (Fig. S25 A-B). The contents of NNN were reduced by 20-60% and 25-60% in all leaf groups in two years. The average content of NNN was diminished to a level less than 0.5 ppm in most leaf groups (Fig. S25 A-B). The contents of NNK were reduced by 5- 50% in all leaf groups in 2011 and 5-60% in four of five leaf groups in 2012. The contents of NAB were reduced by 20-70% and 15-50% in all leaf groups in two years. The contents of NAT were reduced by 50-65-% and 35-70% in all leaf groups in two years. The contents of each NNN, NNK, NAB, and NAT were averaged for all leaf groups of each plant. The resulting data showed that the average contents of each TSNA were significantly reduced by 25-60% (P-value < 0.05) in 2011 and 2012 (Fig. S25 A-B). The contents of four TSNAs were summed for all leaf groups of each plant. The resulting data showed that the total contents of four TSNAs were significantly reduced by 57% and 43% (P-value < 0.05) in 2011 and 2012, respectively (Fig. S26).

## Discussion

Our findings indicate that cis-regulatory elements and TFs are useful molecular tools for a *de novo* design to create a novel regulation of plant secondary metabolism.

Herein, based on the regulation mechanism elucidated (Fig. 2), we constructively term the designed regulation to be a *Distant Pathway-Cross Regulation* (DPCR) of PAP1 and TT8. The rationale is that the pathway of nicotine does not exist in Arabidopsis and two anthocyanin TFs are metabolically characterized to be distant from the tobacco alkaloid pathway. The other reason is that our findings are distinct from the general pleotropic effects that have been generally used to explain gene functions without a mechanistic elucidation. This DPCR definition is characterized by the mechanisms that PAP1, TT8, and their complex formed with WD40 negatively regulate the tobacco alkaloid biosynthesis via activating *NtJAZ* repressor genes. Accordingly, this definition distinguishes from the master regulatory function of PAP1, TT8, and their MBW complex that activates the anthocyanin biosynthesis in Arabidopsis [8, 14, 73, 74] or red tobacco shown here via directly activating most or late pathway genes, such as *DFR*, *ANS*, and *3-GT* (Fig. S10). Based on these data, we propose a model to characterize the DPCR created from the *de novo* design and the regulation of tobacco alkaloid biosynthesis in red tobacco plants (Fig. 8). Given that *NtTTG1*is expressed in roots, leaves, and flowers, this model proposes that PAP1 and TT8 likely anchor to the endogenous tobacco NtTTG1 (WD40) to form a heterogeneous MBW complex, which has a dual regulatory function that regulates tobacco metabolism. First, the MBW complex directly activates the anthocyanin biosynthesis (function I), which is intensively elucidated. Second, the MBW complex upregulates four *NtJAZ1*, *NtJAZ3*, *NtJAZ7*, and *NtJAZ10*, which increase NtJAZ repressors (function II). Next, the increased NtJAZs bind to more MYC2 and block the release of this TF. Then, the reduction of free MYC2 results in downregulating *EFR* and six or seven nicotine pathway genes (Fig. 3). Finally, the biosynthesis of nicotine, OTAs, and four TSNAs is significantly diminished in red tobacco plants (Figs. 4, 6, and 7). In addition, based on the report that AtMYC2 solely regulates the *AtJAZ* transcription via a feed-back regulation [75], this model proposes that PAP1/TT8 and NtMYC2 co-activate the expression of NtJAZs in red tobacco plants. When JA is absent, PAP1 and TT8 play the main regulatory role to activate the expression of four *NtJAZs*. To date, in addition to tobacco, PAP1 and its homologs have been introduced to a few crops to engineer anthocyanins for value-added traits, such as tomato [22, 23], hop [24], canola [25], and rose [76]. Although whether and how PAP1 and TT8 can regulate non-anthocyanin or non-flavonoid pathways in these plants remains unknown, it was interesting that PAP1 transgenic rose plants were reported to produce more than 6.5-time scent terpenoids [76]. Based on our findings, it is interesting to investigate whether a similar DPCR occurs to increase scent terpenoids in rose in the future. Moreover, different TF families are a large part of plant genomes [77–80], our findings suggest that it is necessary to investigate DPCR events of other TFs to fully understand their regulatory functions. In particular, as shown in our data that carcinogenic TSNAs are significantly reduced in red tobacco plants, investigations of potential DPCR of TFs likely create value-added traits for crop improvement and human health.

**Figure 8.**
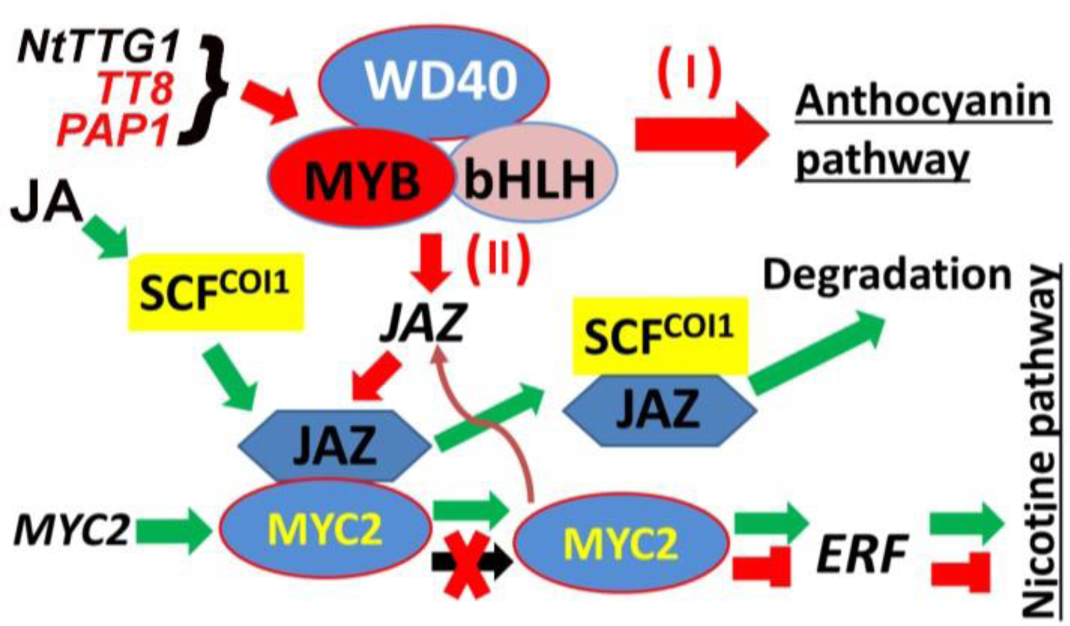
A model showing the distant pathway-cross regulation (DPCR) created for the downregulation of the biosynthesis of tobacco alkaloids in red tobacco plants. Regulation pathways created in red tobacco plants are featured by red colored solid arrows, “X” shape, and “T” blocks. The stacked *PAP1*and *TT8* as well as the endogenous *NtTTG1* encode MYB, bHLH, and WD40 proteins, respectively, which form a complex to regulate two biosynthetic pathways. The regulation pathway (function I) is the activation of biosynthesis of anthocyanin. The regulation pathway (function II) is the DPCR that the MYB-bHLH-WD40 complex upregulates the endogenous *NtJAZs* leading to the reduction of MYC2 and the downregulation of *EAR* and the nicotine biosynthesis. In addition, The JA-JAZ-MYC regulatory pathway is indicated with green arrows starting with JA (greenish arrows). The feedback regulation of NtMYC2 indicated by a brownish bending arrow is proposed to activate the expression of NtJAZs. SCF: Skp/Cullin/F-box and COI1: coronatine insensitive 1.

This *de novo* regulation design with PAP1/TT8 TFs and *NtJAZs* shows a promising application in enhancing efforts for the simultaneous reduction of most harmful tobacco alkaloids. As introduced above, in the past decades, multiple previous studies had been undertaken to develop technologies to reduce harmful tobacco alkaloids, such as gene suppression or silencing technologies with anti-sense or RNAi of pathway genes (Fig. 1 B), including *PMT* [65], *ODC* [64], *QPT* [63], *NDM* [81], A622 [82], *BBL* [49], and others [42]. On the one hand, most studies successfully reported the decrease of the contents of nicotine, OTAs, or TSNAs in tobacco plants grown in the greenhouse, growth chambers, or field. On the other hand, these previous successes revealed a challenge that all tobacco alkaloids could not be diminished simultaneously. For example, the decrease of nicotine led to either the increase of anatabine or non-reduction of other tobacco alkaloids [64, 65]. In addition, the reduction of harmful compounds was relatively limited. For instance, an RNAi of A622 could mainly reduce NNN and total TSNAs [81]. A gene silencing of *BBL* was reported to mainly decrease the nicotine content [49]. Furthermore, all these successes have showed that the reduction of nicotine and other alkaloids to a practical level requires more studies, such as elucidation of metabolic regulation and unknown genes. As intensively characterized, the regulation of the nicotine biosynthesis is via three cohort layers, the presence/absence of an active JA form, the JA and JAZ signaling pathway, and the release of MYC2 for activation of ERF189 and pathway genes [42, 48, 55] (Fig. 1 B). To date, an important unanswered question is how tobacco plants regulate *NtJAZs*. In Arabidopsis, the expression of *AtJAZs* is solely activated by AtMYC2 via a feed-back regulation [75]. Based on this, although NtMYC2 members have not been reported to regulate the transcription of *NtJAZs* in tobacco plants, it can be postulated that NtMYC2 members may also perform a feed- back regulation of *NtJAZs* (Fig. 8). Our *de novo* design with PAP1 and TT8 and MRE and G-box elements was used to target four *NtJAZs*. As designed, PAP1, TT8, PAP1 and TT8 together, and their MBW complex bound and activated promoters of the four NtJAZs (Fig. 2) and the stacked overexpression of *PAP1* and *TT8* led to the upregulation of four *NtJAZs* (Fig. 3 A-B and K-L, Fig. S13). As the positive result, the biosynthesis of nicotine was downregulated in red P+T genotypic plants (Figs. 3 and 4). The contents of nicotine, four OTAs, and four TSNAs were simultaneously reduced in most or all cured leaf groups and entire plant leaves of P-T-NL, P-T-KY, and PAP1 genotypes (Figs. 6 and 7, Figs. S20). The NNN contents were particularly diminished to a level less than 0.55 ppm in most of red tobacco leaves (Fig. 7 A-B).

This low content is significant to human health, because NNN is a severe tobacco carcinogen and its content in any smokeless tobacco products is limited to 1.00 pm proposed by FDA. In addition, other TSNAs were reduced to a level lower than 0.5 ppm in most or all leaves of red P+T genotypic tobacco plants. To further substantiate the function of this *de novo* regulation design, we tested our PAP1 red tobacco variety, in which the expression of *PAP1* activated two *NtTT8* homologs (Fig. S21).

Although this variety was created from Xanthi, a low-nicotine and flue-cured cultivar, the two years’ field trials showed the reduction of nicotine, OTAs, and TSNAs in PAP1 plants (Figs. S23-24 and S25-26). These data indicate that PAP1 can partner with the endogenous tobacco NtTT8 homologs to form a DPCR to downregulate the nicotine biosynthesis. Taken together, these findings provide a new platform to eliminate tobacco carcinogens to a non-harmful level and to alter plant metabolism in other crop plants for novel traits.

The plant kingdom has a large number of TFs that regulate plant development, structure, responses to stresses, and plant metabolisms. In 2000, when the genome of *Arabidopsis thaliana* was sequenced [83], gene annotation and sequence analysis revealed more than 1500 TFs (accounting for more than 5% of annotated genes). Since then, more TFs have been reported from numerous sequenced plant genomes, such as rice [84, 85]and soybean[86], and multiple browser servers have been developed to search TF families and their regulatory functions [87–93]. By 2017, 156 plant genomes were sequenced, including 16 from Chlorophytae, one from Charophyta, one from Marchantiophyta, two from Byrophyta, one from Lycopodiophyta, two from Coniferophyta, one from Basa Magnoliophyta, 38 from monocots, and 95 Eudicots. [67]. These 156 genomes together with sequences of nine additional plant species allowed the annotation of 320,370 plant TFs, which were grouped into 58 families [67]. Likewise, the plant genomes have a large number of binding sites (motifs) (BS), such as G-box and MRE in promoters, to which TFs bind. In recent, a public PlantRegMap server was developed from 63 plant species (out of 156 genome-sequenced plants) to search TFBSs and the interactions between TF and BS [68]. This PlantRegMap characterized a conserved regulatory landscape that included 21,997,501 TFBSs, 21,346 TFs, and more than two million interactions between them [68]. It can be speculated that as more plant species will be sequences, the number of plant TFs will be increased. To our knowledge, none of TFs and BSs has been used as molecular tools for a novel regulatory design to engineer better valuable traits of crops. Based on our first proof-of-concept study with PAP1 vs. MRE and TT8 vs. G-box that are effective molecular tools to design novel regulations to create a DPCR in tobacco plant, it can be speculated that the large number of plant TFs and BSs provide an extremely rich molecular tool source for novel regulatory designs, which will be fundamental for plant engineering.

## Conclusion

The findings show that PAP1, TT8, MREs, and G-box elements are useful molecular tools to design a *De Novo* regulation, a distant pathway-cross regulation (DPCR) of plant secondary metabolism in plants. The findings unearth novel regulatory functions of PAP1 and TT8 that are two positive regulators of *NtJAZs* and negatively regulate tobacco alkaloid biosynthesis in red tobacco plants. This *De Novo* regulation design significantly reduces harmful nicotine, other tobacco alkaloids, and all TSNAs in all or most leaves of tobacco plants. Our findings indicate that the great number of TFs and BSs provide an extremely rich molecular tool source for novel regulatory designs, which will be fundamental in plant engineering for value-added crops and products.

## Supporting information

Supporting materials and growth conditions, Method S1-S8, 26 supporting figures and 9 supporting tables.

## 26 Supporting Figure legends

Figure S1 Promoter sequences of *NtJAZ1* and *NtJAZ3* and identification of cis-elements

Figure S2 Promoter sequences of *NtJAZ7* and *NtJAZ10* and identification of cis-elements

Figure S3 Images showing expression and purification of the binding domains of PAP1 and TT8 induced in *E. coli in vitro*

Figure S4 Promoter sequences of *NtPMT2* and *NtODC* and identification of cis- elements.

Figure S5 Electrophoretic mobility shift assays (EMSAs) showing weak binding of TT8 to G-box-like probe and no binding of PAP1 to MRE-like probe.

Figure S6 Dual-luciferase and Chip-qPCR experiments showing no activation of *NtPMT2* and *NtODC2* promoters bound by PAP1 and TT8 alone, PAP1 and TT8 together, and PAP1-TT8-WD complex.

Figure S7 Diagrams showing gene stacking design for DNA synthesis, binary construct, and selection of T2 homozygous plants.

Figure S8 Comparison of *AtPAP1* and *AtTT8* transgene expressions in roots of wild- type Narrow Leaf Madole (NL) and KY171, P+T transgenic NL and KY171, and vector control transgenic plants.

Figure S9 Phenotypes of wild type, vector-control, and P+T-NL and P+T-KY tobacco plants used for analysis of gene expression and metabolites.

Figure S10 Transcriptional upregulation of eight flavonoid pathway genes in red transgenic P+T-NL and P+T-KY tobacco plants.

Figure S11 Anthocyanin levels in roots and leaves of wild type and red P+T transgenic tobacco plants.

Figure S12 Proanthocyanidin formation in roots of red P+T transgenic tobacco plants Figure S13 Comparison of *NtJAZ7* and *NtJAZ10* gene expression in roots of wild-type Narrow Leaf Madole (NL) and KY171, P+T transgenic NL and KY171, and vector control transgenic plants.

Figure S14 Comparison of *MYC1a*, *MYC1b, MYC2a* and *MYC2b* gene expression in roots of wild-type (WT) Narrow Leaf Madole (NL) and KY171, P+T transgenic NL and KY, and vector control transgenic plants.

Figure S15 Schematic diagram showing procedures for field farming practice and leaf harvest from wild type and red transgenic tobacco plants.

Figure S16 Field design for farming practice of red P+T transgenic plants. Figure S17 Planting, plant growth, and phenotypes of wild type and red P+T transgenic tobacco plants in field from the first day to 75 days after planting. Figure S18 Harvest of plants from field for air-curing and cleanup of field.

Figure S19 Comparison of leaf color after air-curing and grouping of leaves for sampling.

Figure S20 Reduction of total tobacco alkaloids in each leaf group of red P+T-NL1 and P+T-KY1 plants compared to wild type ones.

Figure S21 Upregulation of *NtAn1a* and *NtAn1b* in roots of red PAP1 tobacco plants. Figure S22 Field farming practice of PAP1 tobacco plants and leaf harvest.

Figure S23 Reduction of nicotine, nornicotine, anabasine, and anatabine in PAP1 tobacco leaves.

Figure S24 Reduction of total nicotine, nornicotine, anabasine, and anatabine in all leaves of PAP1 tobacco plant.

Figure S25 Reduction of NNN, NNK, NAB, and NAT in flue-cured leaves of PAP1 tobacco compared to leaves of wild type Xanthi control.

Figure S26 Reduction of total TSNAs in flue-cured leaves of PAP1 tobacco.

## Nine Supporting Table capitals

Table S1 Thermal cycle and sequences of primers designed for PCR, qRT-PCR, QPCR, respectively.

Table S2 Float trays used for seedlings in the water nursery bed in the greenhouse. The same trays were prepared for PAP1 tobacco plants.

Table S3 Instrument Parameters used in Tobacco Specific Nitrosamines analysis. Table S4 HPLC Gradient used in Tobacco Specific Nitrosamines analysis.

Table S5 Typical Parameter Table Triple Quadrupole Mass Spectrometer used in Tobacco Specific Nitrosamines analysis.

Table S6 Reagents and Standards in for Tobacco Specific Nitrosamines analysis. Table S7 Instrument Parameters used in Tobacco Alkaloids analysis.

Table S8 Oven Program used for in Tobacco Alkaloids analysis.

Table S9 Reagents and Standards used for Tobacco Alkaloids analysis.

